# Reduced sera neutralization to Omicron SARS-CoV-2 by both inactivated and protein subunit vaccines and the convalescents

**DOI:** 10.1101/2021.12.16.472391

**Authors:** Xin Zhao, Dedong Li, Wenjing Ruan, Rong Zhang, Anqi Zheng, Shitong Qiao, Xinlei Zheng, Yingze Zhao, Zhihai Chen, Lianpan Dai, Pengcheng Han, George F. Gao

## Abstract

Omicron variant continues to spread all over the world. There are lots of scientific questions remaining to be answered for such a devastating variant. There are a dozen of vaccines already in clinical use. The very urgent scientific question would be whether or not these vaccines can protect Omicron variant. Here, we tested the sera from both convalescents and vaccine recipients receiving either inactivated or protein subunits vaccines (CoronaVac from Sinovac, or BBIBP-CoV from Sinopharm, or ZF2001 from Zhifei longcom) for the binding antibody titers (ELISA) and neutralization antibodies titers (pseudovirus neutralization assay). We showed that Omicron do have severe immune escape in convalescents, with 15 of 16 were negative in neutralization. By contrast, in vaccinees who received three jabs of inactivated or protein subunit vaccine, the neutralizing activity was much better preserved. Especially in the ZF2001 group with an extended period of the second and third jab (4-6 months) remains 100% positive in Omicron neutralization, with only 3.1-folds reduction in neutralizing antibody (NAb) titer. In this case, we proposed that, the multi-boost strategy with an extended interval between the second and third jab for immune maturation would be beneficial for NAb against devastating variants such as Omicron.

---

COVID-19 pandemic continues as a public health threat worldwide^1^. The Omicron variant of concern (VOC), reported on November 24th, already has been detected in 58 countries and regions (WHO). With 32 mutations in its spike (S) protein, including 15 in the receptor binding domain (RBD), the immune escape of Omicron is of highly concern. Inactivated vaccines such as CoronaVac^®^ and BBIBP-CoV^®^, and protein subunit vaccine such as ZF2001^®^ were already widely used in China and several other countries^2^. Until December, 2021, over 8.4 billion doses of vaccines worldwide with more than 2.6 billion doses in China have been injected (https://coronavirus.jhu.edu/).

Here, we analyzed the binding and neutralizing antibodies (NAbs) elicited by three shots of subunit vaccine ZF2001 or three shots of inactivated vaccine (CoronaVac or BBIBP-CoV). Sera from ZF2001 vaccinees were also separated into two groups: one with a shorter interval between the second and third dose (jabs at 0, 1 and 2 months), and the other with a more extended interval (jabs at 0, 1 and around 5 months, Table S1). The inactivated vaccinees all have their third shot more than 6 months after the second one (Table S1). The binding antibody were analyzed by ELISA with trimeric S proteins of either prototype (WH-01) or Omicron. The binding antibody against Omicron has no significant decline in the three-shot inactivated vaccine group (Fig. 1A). However, both ZF2001 groups have significant decrease in binding antibody (Fig. 1B-C).

**Figure 1:**
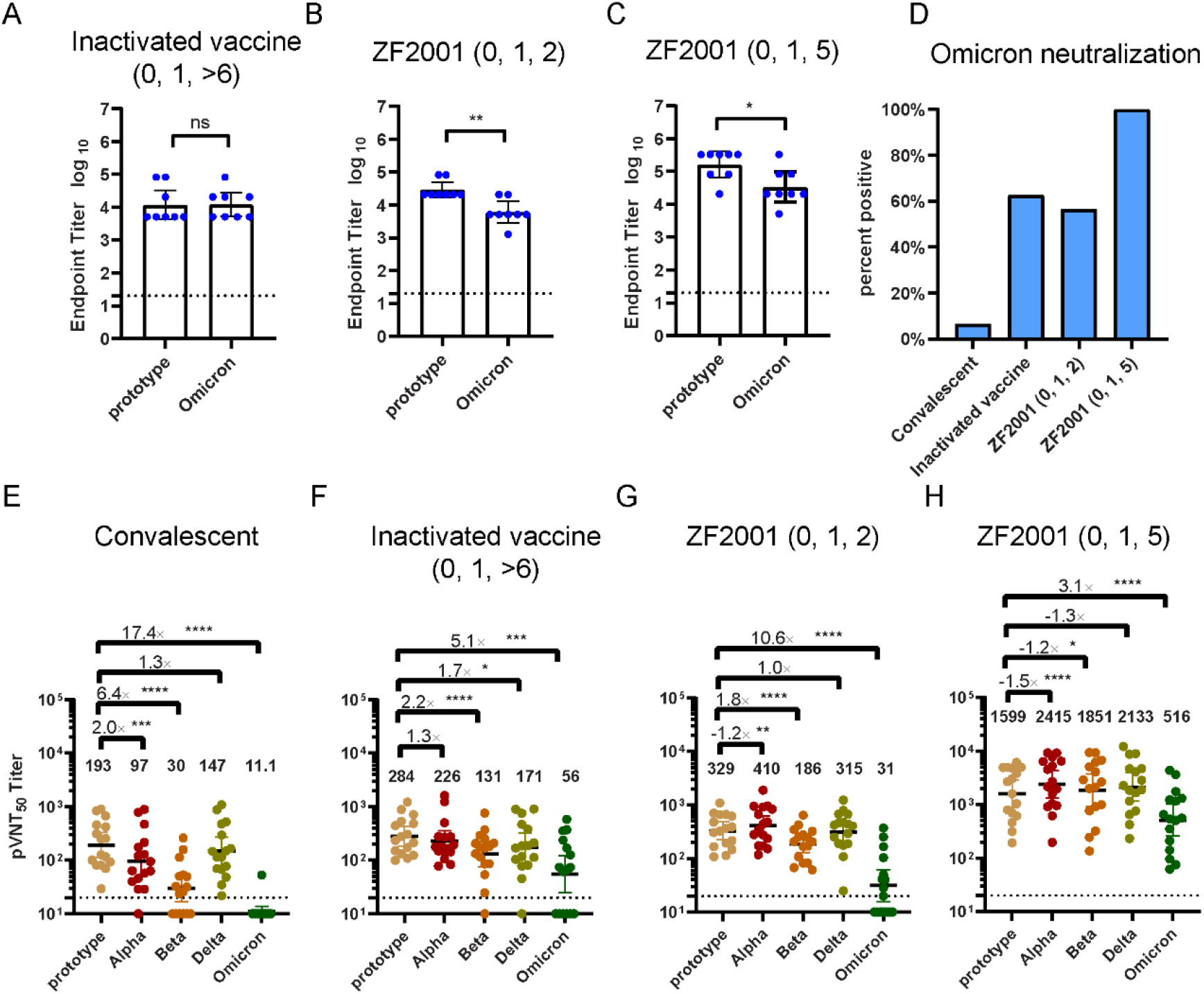
Serum IgG titer and pseudovirus neutralization against Omicron variant. Serum samples from convalescents or vaccinees were collected and the binding and neutralizing antibody (NAb) against SARS-CoV-2 VOCs including Omicron were detected. **Panel A-C** show the serum IgG titer tested by ELISA against SARS-CoV-2 prototype or Omicron trimeric spike (S) protein in the three-shot inactivated vaccine group (**A**), ZF2001 (0, 1, 2) group (**B**), and ZF2001 (0, 1, 5) group (**C**). The statistical significance were analyzed by Mann-Whitney test (n=8 each, *, p<0.05; **, p<0.01; ***, p<0.001 and ****, p<0.0001). **Panel D** shows the percentage of samples tested positive (>1:20) in Omicron neutralization. **Panel E-H** shows the 50% pseudovirus neutralization (pVNT_50_) of sera against SARS-CoV-2 prototype and VOCs (Alpha, Beta, Delta and Omicron). Sera from convalescents (**E**, n=16), vaccinees taken three shot of inactivated vaccine (**F**, n=16), ZF2001 protein subunit vaccine in a 0, 1, 2 regimen (**G**, n=16) or 0, 1, 5 regimen (**H**, n=16) were tested. Geometric mean titer (GMT) was indicated on each column. The statistical significance were analyzed by two-tailed Wilcoxon matched-pairs signed-rank test (*, p<0.05; **, p<0.01; ***, p<0.001 and ****, p<0.0001). All neutralization assays were repeated twice. GMT with 95% confidential interval (CI) were shown for all data. The lower detection limit (1:20) was marked by a dashed line. GMTs lower than 1:20 were indicated as IgG or neutralization antibody negative and counted as 10 in statistical analysis.

Further, we tested the NAbs of the serum samples against SARS-CoV-2 prototype and VOCs including Omicron by pseudovirus neutralization assay (Fig 1D-H and Fig S1). 15 of 16 convalescent samples were detected negative in Omicron NAbs (Fig. 1D), indicating that the immune escape of Omicron is sever, which is in accordance with other recent unpublished serological and monoclonal antibody analyses^3^. However, people have three shots of vaccines mostly preserved the neutralization against Omicron. 62.5% persons taken three shots of inactivated vaccines, 56.25% ZF2001 (0, 1, 2 regimen) and 100% ZF2001 (0, 1, 5 regimen) vaccinees are Omicron NAb positive (Fig. 1D). As for the NAb titer, the convalescent group have a 17.4 times reduction from the prototype to Omicron (Fig. 1E). The inactivated vaccine group have 5.1 times reduction (Fig. 1F), the ZF2001 (0, 1, 2) groups have a 10.6 times reduction; and the ZF2001 (0, 1, 5) group have only a 3 times reduction in NAb titers against Omicron (Fig. 1G-H). Moreover, as we reported^4^, the longer interval between the second and third shot can elicit higher antibody titers and better neutralization against all variants.

The findings prompt us to consider that multiple boost and prolonged intervals for vaccinations might be beneficial to conquer the seriously mutated variants such as Omicron. This is in accordance with the researches on mRNA vaccines which havereported that people with three shots much better preserved the neutralization against Omicron variant (^5^ and BioNtech reports). In this case, a booster shot should be suggested for people who have received two shots or have been previously infected by SARS-CoV-2. For vaccines such as ZF2001 which already have a three shots regimen, a prolonged interval between the second and the third shot would be beneficial. The next generation vaccines with a broad protection are also needed. However, more real-world data are needed for a precise judgement.

