## supplementary for "Reduced sera neutralization to Omicron SARS-CoV-2 by both inactivated and protein subunit vaccines and the convalescents"

### Supplementary Appendix

**Contents** | **Page**

#### Materials and Methods

##### Serum samples

The serum sample of convalescents were provided by Ditan Hospital, Beijing, China. The study was approved by the Ethics Committee of the Institute of Microbiology, Chinese Academy of Sciences (SQIMCAS2021149) and National Institute for Viral Disease Control and Prevention, Chinese Center for Disease Control and Prevention (IVDC2020-021). All candidates signed the written informed consent.

According to vaccination strategies, participants are classified into four groups with 16 each: i) convalescents 4-5 weeks after tested negative in SARS-CoV-2; ii) 3 doses of inactivated vaccines (CoronaVac or BBIBP-CorV, 8 samples each); iii) 3 doses of recombinant protein subunit vaccine (ZF001) with an interval of 1 month between each vaccination; iv) 3 doses of ZF001 with an extended interval between the second and the third dose for approximately 5 month. For convalescents, only NAb positive samples against prototype was picked. And for the other three vaccinated groups, samples were randomly picked. Detailed information are available in **Table S1**.

##### ELISA

ELISA were performed as mentioned in our previous work[1]. Briefly, the 96 well ELISA Plates (Corning Incorporated， USA) were coated at 4℃ overnight with 100 ng SARS-CoV-2 prototype trimeric S protein (SPN-C52H9, Acro, China) or SARS-CoV-2 B.1.1.529/Omicron trimeric S protein (SPN-C52Hz, Acro, China). After 1 h blocking using 5% defatted milk, serum samples were added at a 4-time serial dilution starting from 1:20. Plates were subsequently incubated with goat anti-human IgG (H&L), HRP Conjugated (BE0122, EASYBIO, China) and TMB (P0209-500ml, Beyotime, China) substrate. Each plate included negative control. Absorbance greater than 2.5-fold of the background values was considered positive.

##### Pseudotyped virus neutralization assay

The construction of VSV-ΔG-GFP based SARS-CoV-2 pseudotyped virus was mentioned in previous work with slight modifications[2,3]. pCAGGS vector was used to construct recombinant plasmid of codon optimized spike proteins of SARS-CoV-2 wild type (Wuhan-1 reference strain) and variants, with an 18 amino acid truncation at the C-terminal of the spike protein (with mutations shown in **Fig. S1**). 30 μg of the construct was transfected into HEK 293T cells respectively to prepare corresponding packaging cell lines of each strain. VSV-ΔG-G-GFP pseudovirus were added 24 h after the transfection. After 1 h infection, VSV-ΔG-G-GFP was removed by changing the medium into fresh complete DMEM medium with anti-VSV-G antibody (I1HybridomaATCC® CRL2700™). Supernatants were collected after another 30 h incubation, passed through a 0.45 μm filter (Millipore, Cat#SLHP033RB), aliquoted and stored at -80 °C.

For the neutralization assay, the heat-inactivated (56 °C for 30min) serum samples were 2-fold serial diluted starting from 1:20. Equivalent pseudovirus (1000 transducing units, TU) were incubated with the sera at 37 °C for 1 h, and the mixture was then added onto pre-plated Vero cells (ATCC CCL81) in 96 well plate. The TU numbers were calculated on a CQ1 confocal image cytometer (Yokogawa) after a 15 h incubation[2].

**Data analyzing and statistical analysis**

Software provided with CQ1 confocal image cytometer was used to calculate the TU numbers of neutralization assay and the statistical analyses were performed by GraphPad Prism 8.0 (GraphPad Software Inc.). $pVNT_{50}$ were determined by a non-linear regression. $pVNT_{50}$ below the lower limit of detection (<20) was recorded as 10 in the geometric mean calculation. Neutralization between prototype and different variants were compared using a two-tailed Wilcoxon matched-pairs signed rank test.

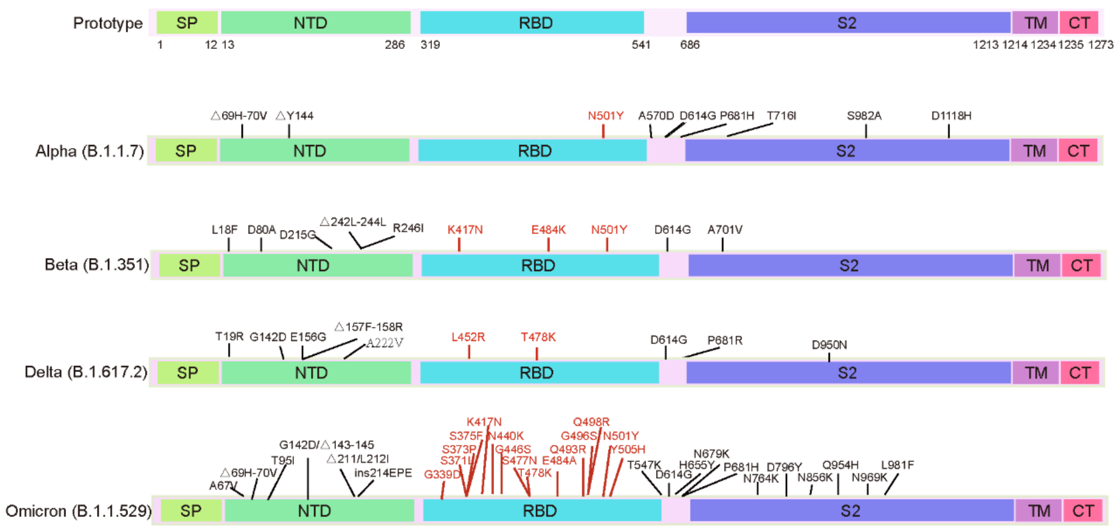

**Figure S1: Schematic representation of the mutations on SARS-CoV-2 Spike proteins.** The mutation sites of the variants were marked on top of the spike protein of each variant. The mutations on receptor binding domain were colored in red. (SP: signal peptide, NTD: N-terminal domain, RBD: receptor-binding domain, TM: transmembrane domain, CT: C-terminal cytoplasmic domain).

**Table S1. Characteristics of vaccine recipients.**

| Groups | ZF2001 (0, 1, 2) | ZF2001 (0, 1, 5) | Inactivated vaccine (0, 1, >6) |
|---|---|---|---|
| **No.of participants** | 16 | 16 | 16 |
| **Age (median, range)** | 29 (24-54) | 30 (23-52) | 25.5 (23-38) |
| **Sex** | | | |
| Male (%) | 4 (25.0) | 6 (37.5) | 9 (56.3) |
| Female (%) | 12 (75.0) | 10 (62.5) | 7 (43.7) |
| **Time interval between the first and second doses (Days, Median, range)** | 29 (28-35) | 28 (27-35) | 26.5 (15-39) |
| **Time interval between the second and third doses (Days, Median, range)** | 35 (31-47) | 118 (82-309) | 219.5 (188-459) |
| **Time interval between the third dose and blood sampling (Days, Median, range)** | 14 (14-14) | 22 (22-73) | 26.5 (8-39) |

#### Acknowledgments

We thank Beijing Ditan hospital for providing us the convalescents' serum samples. We thank all the volunteers for providing blood samples. This work was supported by the National Key R&D Program of China (2020YFA0907102 and 2020YFA0509202). The Strategic Priority Research Program of the Chinese Academy of Sciences (XDB29010202) and the intramural special grant for SARS-CoV-2 research from the Chinese Academy of Sciences to G.F.G. This work is supported by the National Natural Science Foundation of China, China (NSFC) (81991494), a grant from the Bill & Melinda Gates Foundation (INV-006377). X.Z. is supported by Beijing Nova program of Science and Technology (Z191100001119030), and Youth Innovation Promotion Association of the CAS (20200920). L.D. is supported by the Excellent Young Scientist Program from NSFC (82122031) and Youth Innovation Promotion Association of the CAS (2018113).

#### Author contributions

GFG conceived and designed the study. GFG, XZ, PH and LD designed, and coordinated the experiments. XZ, DL, WR, RZ, AZ, SQ, XZ and YZ performed experiments. XZ and RZ analyzed the data. ZC, YZ, DL, and WR helped in the sample collection and organization. GFG, XZ, LD, PH and XZ drafted and revised the manuscript. All authors reviewed and approved the final manuscript.

#### Declaration of interests

L.D. and G.F.G. are listed in the patent as the inventors of the RBD-dimer as a betacoronavirus vaccine. The patent has been licensed to Anhui Zhifei Longcom for protein subunit COVID-19 vaccine development. All other authors declare no competing interests.
